# Electrical spiking of psilocybin fungi

**DOI:** 10.1101/2022.07.02.498545

**Authors:** Antoni Gandia, Andrew Adamatzky

## Abstract

Psilocybin fungi, aka “magic” mushrooms, are well known for inducing colourful and visionary states of mind. Such psychoactive properties and the ease of cultivating their basidiocarps within low-tech setups make psilocybin fungi promising pharmacological tools for mental health applications. Understanding of the intrinsic electrical patterns occurring during the mycelial growth can be utilised for better monitoring the physiological states and needs of these species. In this study we aimed to shed light on this matter by characterising the extra-cellular electrical potential of two popular species of psilocybin fungi: *Psilocybe tampanensis* and *P. cubensis*. As in previous experiments with other common edible mushrooms, the undisturbed fungi have shown to generate electric potential spikes and trains of spiking activity. This short analysis provides a proof of intrinsic electrical communication in psilocybin fungi, and further establishes these fungi as a valuable tool for studying fungal electro-physiology.

## 1. Introduction

Psilocybin fungi, popularly known as “magic” mushrooms, are a group of different species of psychoactive basidiomycetes that have gained an immense popularity since the ethnomycologists Gordon Wasson and his wife Valentina Pavlovna Wasson introduced them to the western audiences in 1957 [1, 2, 3]. Psilocybin fungi are remarkably famous for inducing mystical-type experiences thanks to tryptamine alkaloids contained in its hyphae, mainly psilocybin, baeocystin and norbaeocystin [4, 5], to which the community of users and a growing pool of scientific evidence grants different potential benefits, such as treating depression to helping manage alcoholism and drug addiction [6, 7, 8, 9, 10, 11].

These organisms have been ever since surrounded of an aura of mysticism and criticism in equal shares, with opinions mostly tied to religious or political beliefs rather than being based in scientific research. Nevertheless, magic mushroom have been used by different human cultures across the globe for millennia, probably since the dawn of mankind, as a tool for exploring and healing psychological and physical disorders, or simply to inspire awe, creativity, introspection, and a better appreciation for nature [12, 13, 14, 15, 16, 17, 18]. Considering their cultural and psycho-pharmaceutical importance, the scientific community is trying to make sense of different aspects of their ecology, physiology, pharmacology, and overall, potential biotechnological applications favouring human society and Earth’s biosphere.

Recent research suggests that spontaneous electrical low-frequency oscillations (SELFOs) are found across most organisms on Earth, from bacteria to humans, including fungi, playing an important role as electrical organisation signals that guide the development of an organism [19]. Considering its potential function as communication and integration waves, detecting and translating SELFOs in psilocybin fungi species may result of great utility in understanding the growth and behaviour of these organisms, a knowledge that could be added to the toolbox of cultivation and pharmacological optimisation techniques used by the fungal biotech industry.

Thereby, we recorded the extracellular electrical potential in mushrooms and mycelium-colonised substrates as indicators of the fungi intrinsic activity. Action potential-like spikes of electrical potential have been observed using intra-cellular recording of mycelium of *Neurospora crassa* [20] and further confirmed in intra-cellular recordings of action potential in hyphae of *Pleurotus ostreatus* and *Armillaria gallica* [21] and in extra-cellular recordings of basidiocarps of and substrates colonised by mycelium of *P. ostreatus* [22], *Ganoderma resinaceum* [23], and *Omphalotus nidiformis, Flammulina velutipes, Schizophyllum commune* and *Cordyceps militaris* [24]. While the exact nature of the travelling spikes remains uncertain we can speculate, by drawing analogies with oscillations of electrical potential of slime mould *Physarum poly-cephalum* [25, 26, 27, 28], that the spikes in fungi are triggered by calcium waves, reversing of cytoplasmic flow, and translocation of nutrients and metabolites.

The paper is structured as follows. First, we present the experimental setup in Sect. 2. Secondly, in Sect. 3 we analysed the electrical activity of the fungi. Finally, section 4 presents some conclusions and directions for further research.

## 2. Methods

Two widely distributed species of psilocybin fungi, namely Psilocybe cubensis strain “B+” (Mondo Mycologicals BV, NL), and *Psilocybe tampanensis* strain “ATL7” (Mimosa Therapeutics BV, NL), were cultured separately on a mixture of hemp shavings amended with 5% wheat flour, at 60% moisture content, in polypropylene (PP5) filter-patch bags.^1^ Electrical activity of the basidiocarps and the colonised substrate was recorded using pairs of iridium-coated stainless steel sub-dermal needle electrodes (Spes Medica S.r.l., Italy), with twisted cables and ADC-24 (Pico Technology, UK) high-resolution data logger with a 24-bit analog-to-digital converter (Fig. 1). We recorded electrical activity at one sample per second. During the recording, the logger has been doing as many measurements as possible (typically up to 600 per second) and saving the average value. We set the acquisition voltage range to 156 mV with an offset accuracy of 9 *µ*V at 1 Hz to maintain a gain error of 0.1%. Each electrode pair was considered independent with the noise-free resolution of 17 bits and conversion time of 60 ms. Each pair of electrodes, called channels, reported a difference of the electrical potential between the electrodes. Distance between electrodes was 1-2 cm. We have conducted eight experiments, in each experiments we recorded electrical activity of the fungi via four channels, i.e. 32 recordings in total.

**Figure 1:**
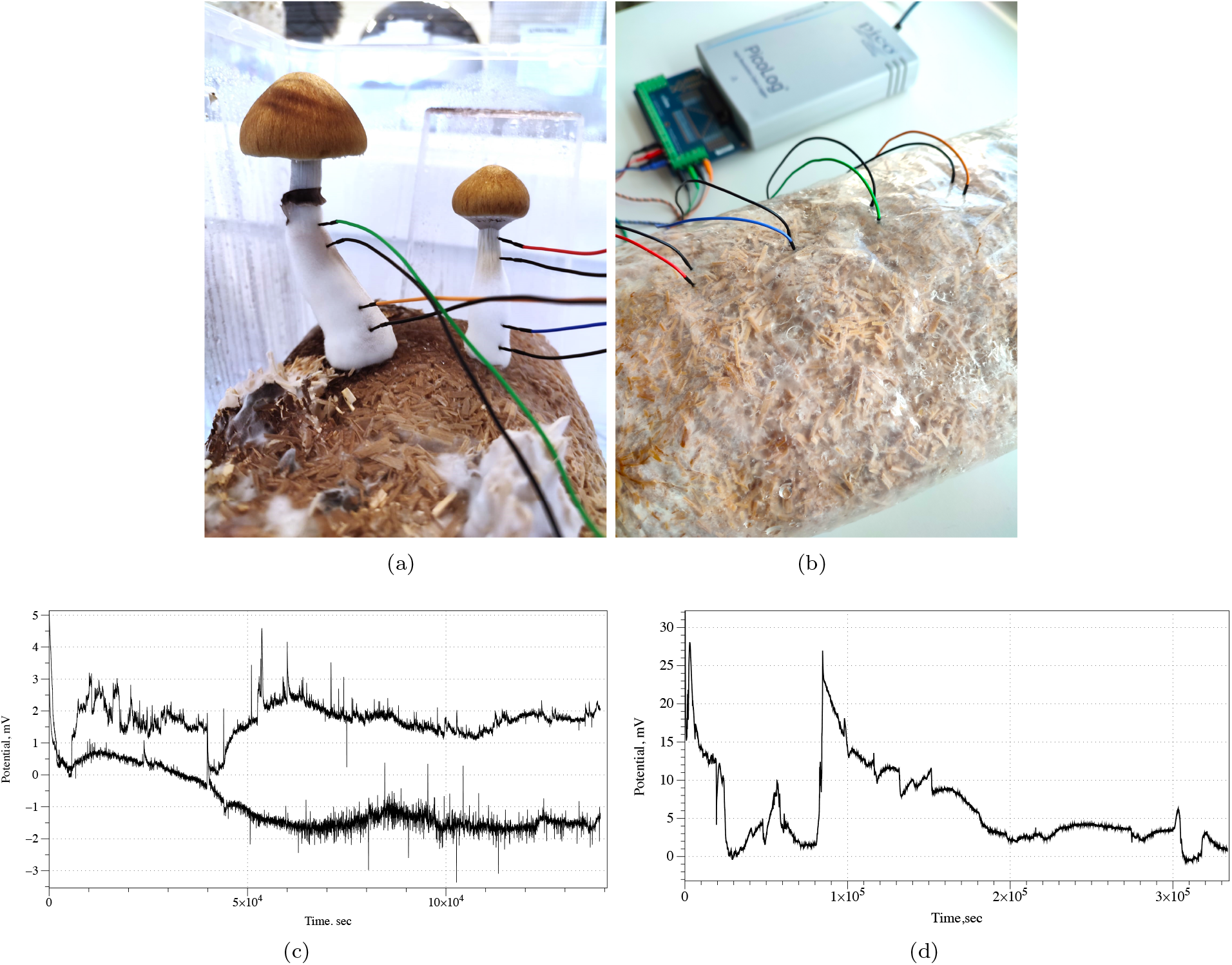
Experimental setup. (a) Example of recording from Psilocybe cubensis basidiocarps. (b) Example of recording from Psilocybe tampanensis mycelium-colonised substrate and view of the experimental setup, in which the electrodes with cables and Pico ADC-24 are seen. (cd) Examples of electrical activity of (c) Psilocybe cubensis, two channels, and (d) Psilocybe tampanensis, one channel.

## 3. Results

Electrical activity recorded from both species of psilocybin fungi shows a rich dynamics of electrical potential. Examples of the recording conducted for nearly four days are show in Fig. 1de. Drift of the base potential can be up to 10-15 mV however rate of the base potential change is measured in days therefore it does affect our ability to recognise spikes. Plots also show that in some cases the signal-to-noise ratio might be substantially low. We omitted such cases from the spike detection pool.

We observed action-potential like spikes of electrical potential. Most expressive spikes, see e.g. Fig. 2a, shown very characteristics of action potential recorded in nervous system with distinctive depolarisation and repolarisation phases and a refractory period. In the exemplar action-potential like spike shown in Fig. 2a depolarisation phase is c. 18 sec up to 4.5 mV; re-polarisation phase is 97 sec; refractory period is rather long c. 450 sec.

**Figure 2:**
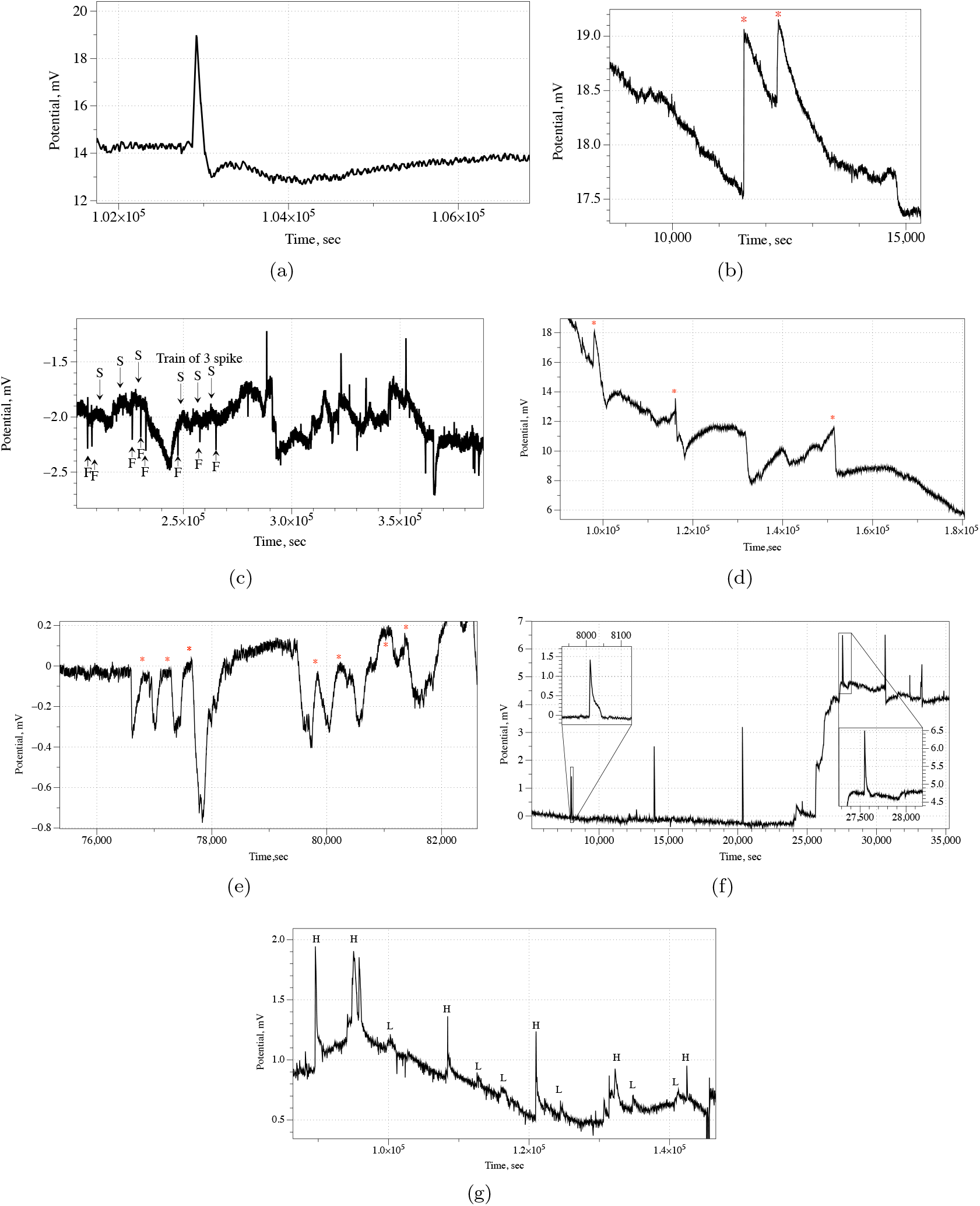
(a) Example of an action-potential like voltage spike, Psilocybe cubensis. (b) Train of two spikes, Psilocybe cubensis (c) Example of fast and slow spike activity,Psilocybe cubensis. An average duration of a fast spike is 3 min. An average duration of a slow spike is 16 min. Examples of fast spikes are labelled ‘F’ and slow ‘S’. Train of three slow spikes is also marked. (d) Example of three spikes in electrical potential of Psilocybe tampanensis, peaks of the spike are labelled ⋆. (e) Two trains of spikes recorded in Psilocybe cubensis: one train comprises of three spikes, another of four spikes; spike are marked by ⋆. (f) Very fast, average 1.5 min, spikes of electrical potential recorded in Psilocybe cubensis. (g) Co-existence of high amplitude, labelled ‘H’, and low amplitude, labelled ‘L’, spikes in electrical activity of Psilocybe cubensis.

In some cases, as illustrated in Fig. 2b, two action-potential like spikes can occur at so short interval that they almost merge. In this particular example, an average spike duration is 13 min, and average amplitude is 1.4 mV.

More commonly the spikes emerge in the trains of spikes. A train is a sequence of spikes where distance between two consecutive spikes does not exceed an average duration of a spike. Two trains of spikes are shown in Fig. 2e. Also, spike can stand alone, as shown in Fig. 2d.

Amongst many types of spike classed by their duration we can select very fast spikes, with duration of 1-2 min, and slow spikes, which width can be 15-60 min. An example of very fast spikes directly co-existing with slow spikes is shown in Fig. 2c. In some cases, only very fast spikes can be observed during the whole duration of the recording, see an example in Fig. 2f.

A co-existence of spikes with high, 0.5-1 mV, and low, 0.1-0.3 mV, is evidence in the recording plotted in Fig. 2g. An amplitude, however, might be not a good characteristic of spikes because it only indicated how far away a wave-front of propagating electrical activity was from a pair of differential electrodes.

Distribution of spike amplitudes version spike width is shown in Fig. 3a. Pearson correlation *R* = 0.0753 calculated on the distribution is technically a positive correlation, however it is low value shows that the relationship between spike width and amplitude is weak.

**Figure 3:**
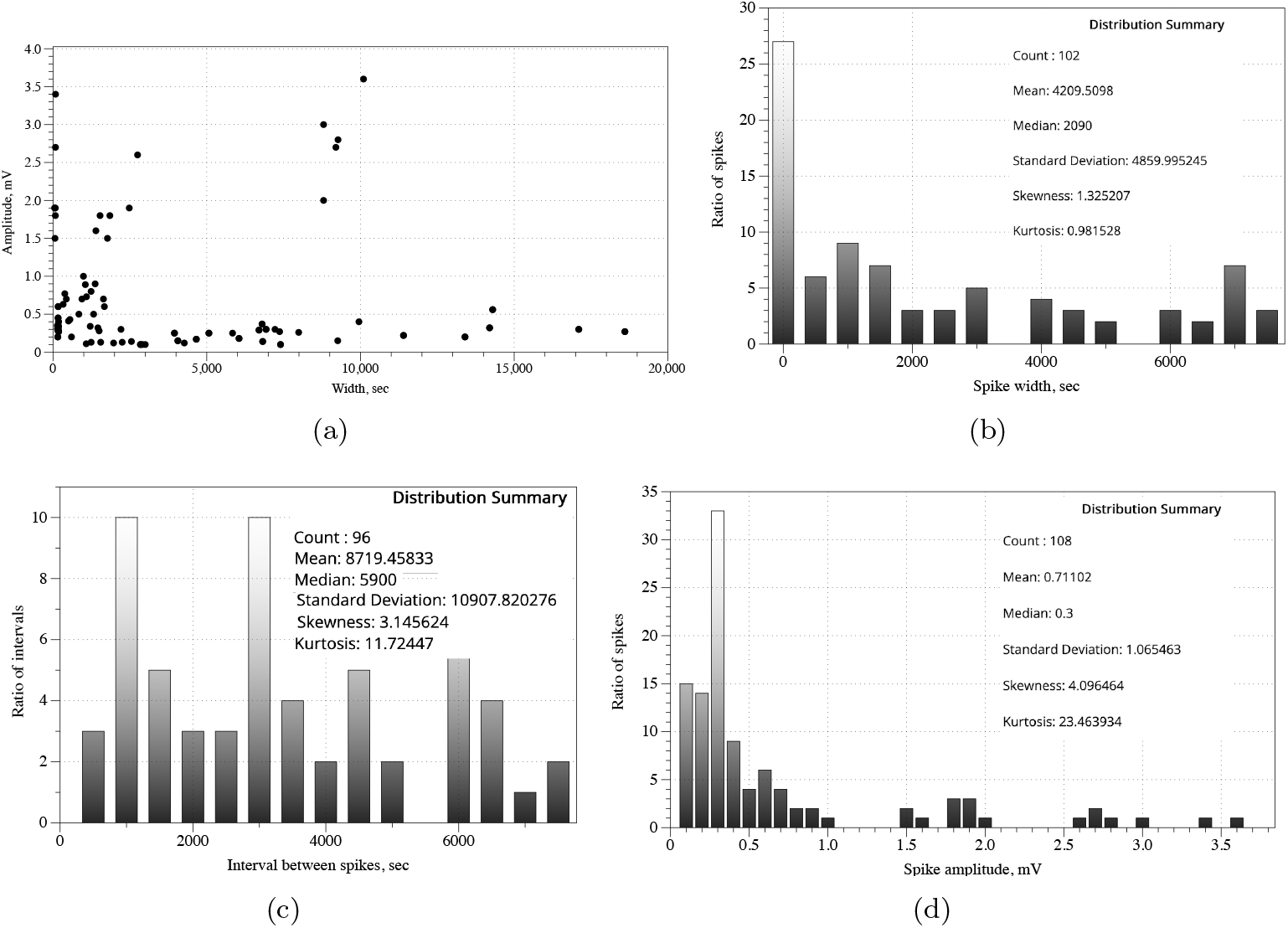
(a) Spike width versus spike amplitude distribution constructed on recording from bother species of fungi studied. (b) Distribution of spike widths, (c) Distribution of interval between spikes. Bin size is 500 in both distributions. (d) Distribution of spike amplitudes, bin size is 0.1.

Distributions of spike width (Fig. 3b), intervals between spikes (Fig. 3c) and spike amplitudes (Fig. 3d) are not normal. This is demonstrated by Kolmogorov-Smirnov test of normality. Values of Kolmogorov-Smirnov statistic is 0.19467 for width distribution, 0.21914 for interval distribution, and 0.28243 for amplitude distribution. Corresponding p-values are 0.00072, 0.00016 and 0.00001.

Integrative parameters of spiking behaviour are the following (Tab. 1). Average duration of a spike is 70 min (*σ* =81 min), median duration is 35 min. Average amplitude of a spike is 0.71 mV (*σ* =1.06 mV), median is 0.3 mV. Average distance between spikes is 145 min (*σ* =181 min), median is 98 min. That is average/median distance between two spikes is a double of the average/media duration of a spike. By the definition of the train, this means that most spikes observed are solitary spikes. Standard deviations of spike duration and amplitude and of interval between spikes are higher thank respective average values. This indicates data are more spread out and we should look out for distinct families of spikes.

Let us separate species and — if any – families of spikes in each species. In *Psilocybe tampanensis* spikes are relatively uniform (Tab. 1): average duration 104 min with *σ* =69 min, average distance between spikes is over three hours and median distance equal to average. In *Psilocybe cubensis* we propose three families of spikes: fast spike, up to 3 min duration, slow spikes, up to 6 hr duration, and very slow spikes, up to 2 days (Tab. 1). Fast spikes rarely form train, an average distance between fast spikes is 17 hr. An average duration of a slow spike is 3.9 hr with an average distance between slow spikes is 9 hr. Average amplitudes of fast and slow spikes are comparable, 0.72 mV and 0.68 mV, respectively. An average duration of a very slow spike of *Psilocybe cubensis* is 22 hr with an average distance between spikes of 33 hr. The very slow spikes have, comparatively to fast and slow spikes, low amplitude of 0.24 mV in average.

**Table 1:** Statistical parameters of spiking. In each cell we show average, standard deviation and median.

| Species | Duration, sec | Amplitude, mV | Distance, sec |
| --- | --- | --- | --- |
| Over all species | 4209, 4859, 2090 | 0.71, 1.06, 0.3 | 8719, 10907, 5900 |
| *Psilocybe tampanensis* | 6246, 4196, 8800 | 2.33, 2.22, 1.71 | 17566, 11183, 17300 |
| *Psilocybe cubensis* | 4082, 4892, 1640 | 0.48, 0.56, 0.30 | 8000, 10458, 4930 |
| *P. cubensis*, fast spikes (1-3 min) | 148, 38, 165 | 0.72, 0.83, 0.35 | 6128, 4516, 3955 |
| *P. cubensis*, slow spikes (up to 6 hr duration) | 1408, 274, 1394 | 0.68, 0.60, 0.50 | 3243, 2125, 3105 |
| *P. cubensis*, very slow spikes (up to 2 days) | 7943, 4658, 6810 | 0.24, 0.11, 0.25 | 11948, 13639, 7880 |

## 4. Discussion

We found that psilocybin fungi exhibit a rich spectrum of oscillations of extracellular electrical potential. We illustrated several types of oscillations and characterised families of fast, slow and very slow oscillations. We demonstrated that several scales — minutes, hours and day — of oscillators states co-exist in basidiocarps and mycelium network of psilocybin fungi. This co-existence is similar to electrical oscillation of a human brain, where fast oscillations might be related to responses to stimulation, including endogenous stimulation by release of nutrients, and slow oscillations might be responsible for memory consolidation [29, 30, 31, 32]. Future research could be concerned with decoding and understanding the spiking events to monitor growth, development and physiological states and overall condition of the fungi both in cultivation setups and natural environments. If we were able to decode spiking patterns of fungi we would be able of ‘speaking back’ to the mycelial network to manipulate the network’s morphology, behaviour and, potentially, enhance production of basidiocarps and sclerotia.

## Acknowledgement

This project has received funding from the European Union’s Horizon 2020 research and innovation programme FET OPEN “Challenging current thinking” under grant agreement No 858132.

## Footnotes

[1] Experiments were conducted at Mimosa Therapeutics BV, The Netherlands.

